# Harmonic-Balance Circuit Analysis for Electro-Neural Interfaces

**DOI:** 10.1101/2020.01.17.910968

**Authors:** Zhijie Charles Chen, Bingyi Wang, Daniel Palanker

## Abstract

**Objective:** Avoidance of the adverse electrochemical reactions at the electrode-electrolyte interface defines the voltage safety window and limits the charge injection capacity (CIC) of an electrode material. For an electrode that is not ideally capacitive, the CIC depends on the waveform of the stimulus. We study the modeling of the charge injection dynamics to optimize the waveforms for efficient neural stimulation within the electrochemical safety limits.

**Approach:** The charge injection dynamics at the electrode-electrolyte interface is typically characterized by the electrochemical impedance spectrum, and is often approximated by discrete-element circuit models. We compare the modeling of the complete circuit, including a non-linear driver such as a photodiode, based on the harmonic-balance (HB) analysis with the analysis based on various discrete element approximations. To validate the modeling results, we performed experiments with iridium-oxide electrodes driven by a current source with diodes in parallel, which mimics a photovoltaic circuit.

**Main results:** Application of HB analysis based on a full impedance spectrum in frequency domain eliminates the complication of finding the discrete-element circuit model in traditional approaches. HB-based results agree with the experimental data better than the discrete-element circuit analysis. HB technique can be applied not only to demonstrate the circuit response to periodic stimulation, but also to describe the initial transient behavior when a burst waveform is applied.

**Significance:** HB-based circuit analysis accurately describes the dynamics of electrode-electrolyte interfaces and driving circuits for all pulsing schemes. This allows optimizing the stimulus waveform to maximize the CIC, based on the impedance spectrum alone.

## 1 Introduction

Electrical stimulation of neurons requires injection of a sufficient amount of charge through the electrode-electrolyte interface. The stimulation threshold depends on various factors, including the electrode geometry [1,2], proximity to the neurons [3], the pulse waveform [2,4], the pulse duration [5, 6], and the repetition rate [2, 7]. When too much charge is injected, the potential drop across the electrode-electrolyte interface would rise beyond the safety window, enabling adverse electrochemical reactions, such as water electrolysis, which may result in damage to the electrode material and biological tissues. The charge injection capacity (CIC) of an electrode is defined as the maximum amount of charge that can be injected in one pulse without driving the electrode-electrolyte interface beyond the safety window [8]. For the efficacy of neural stimulation, bio-compatible electrode materials with large CIC are desirable. Iridium oxides (IrOx) [9–14], platinum black (Pt-Black) [15] and poly(3,4-ethylene-dioxythiophene)-poly(styrenesulfonate) (PEDOT:PSS) [7, 16–18] are some common materials used as neural electrodes for this reason.

The CIC is related to the material capacitance and is affected by the impedance spectrum over a frequency range corresponding to the applied stimulus waveform [11, 19]. Very often, however, the impedance is reported only at one specific frequency, 1 kHz, for example [20], or even not reported at all [7, 13, 14, 17, 18]. Since highly capacitive materials are usually porous and often involve reversible electrochemical reactions, the electro-electrolyte interface is not ideally capacitive. Therefore, dynamics of the charge injection and the CIC depend on the applied waveform. For example, the impedance spectrum of sputtered iridium oxide film (SIROF) shows phase characteristics of the transmission lines, due to the pore resistance and the mass-transfer limitation associated with its porous structure [11, 21]. Hence, the CIC was found to vary with pulse duration [22]. Therefore, materials cannot be characterized by a single CIC value, but should rather be described in a more comprehensive manner, taking into account the impedance spectrum and the waveform of stimulus.

Traditionally, electrode-electrolyte interfaces are described by circuit models with discrete elements, the values of which are fit to the impedance spectrum. Solving the differential equations corresponding to such circuits should describe the resulting charge injection dynamics [23–25]. However, as we will show in Section 3.2, depending on the applied waveform, various discrete-element approximations to the impedance spectrum may result in highly variable outcomes. An accurate representation of the impedance spectrum typically involves constant phase elements (CPEs) [26], the dynamics of which needs to be modeled in frequency domain through the Laplace transform [27], causing extra complications and often incompatible with nonlinear circuit elements, such as diodes, for example.

In this paper, we propose a framework to describe the circuit response without the need to fit an equivalent discrete-element circuit model. If the circuit is driven by controlled current or voltage, directly applying the impedance spectrum in frequency domain yields the corresponding voltage or current response. In cases where neither current nor voltage is directly controlled, we use harmonic-balance (HB) analysis to solve for the response when the circuit driver is nonlinear. For example, photovoltaic retinal prosthesis is driven by photodiodes, which have nonlinear relationship between the voltage and the current [10]. The results of our modeling are validated by comparing with the actual experimental measurements and with the time-domain solutions of the discrete-element circuit approximation. Since the wireless photovoltaic prosthesis is inaccessible to direct measurements in vivo, we developed an experimental technique to mimic the prosthetic circuit without using light. The proposed framework offers a powerful tool for optimizing the design of electro-neural interfaces in general, and predicts, among other useful metrics, the CIC under any stimulation waveform for any electrode material.

## 2 Methods

### 2.1 Circuit Modeling

The circuit for neural stimulation generally consists of two parts: the circuit driver and the load, as shown in Figure 1. The circuit driver is a controlled power source, while the load comprises an electrochemical cell and other power consuming circuit components. These two parts are connected at a port. The output current of the circuit driver is a function of time *t*, denoted by *I*_*D*_(*t*). Similarly, *I*_*L*_(*t*) represents the current flowing through the load, and *V* (*t*) the voltage at the port. The polarities of *I*_*D*_, *I*_*L*_ and *V* are defined as in Figure 1. By Kirchhoff’s laws, for all *t*, we have

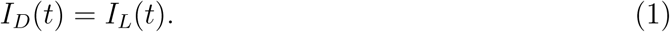

**Figure 1:**
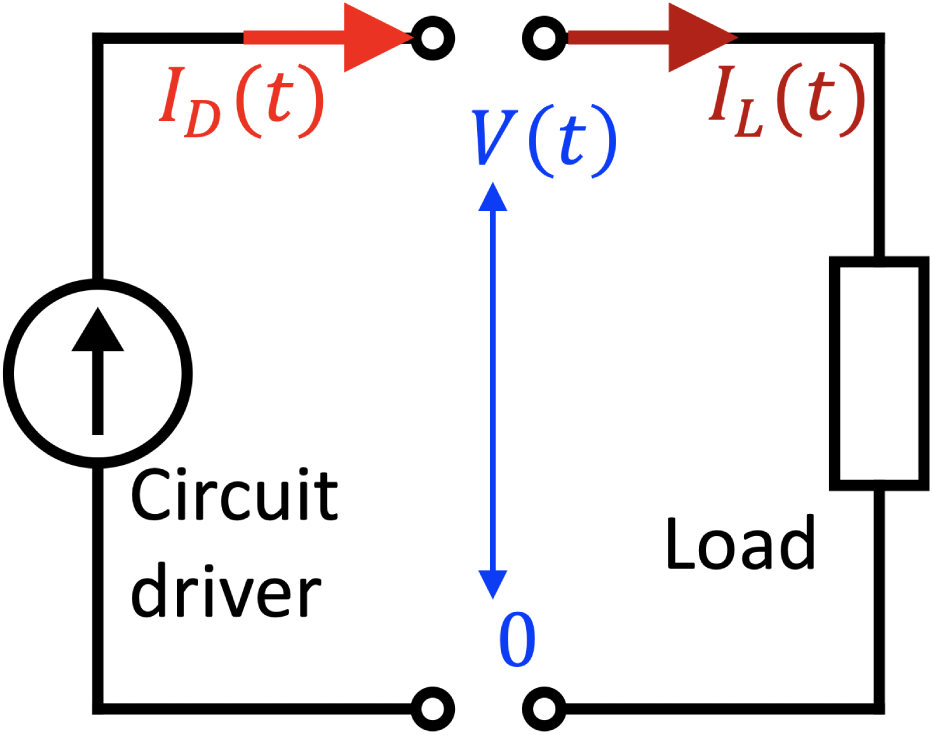
The schematic diagram of the circuit model consisting of the driver and the load, connected by a port. *I*_*D*_(*t*), *I*_*L*_(*t*) and *V* (*t*) are the current output of the circuit driver, the current flowing through the load and the voltage at the port, respectively.

The load is typically a lumped circuit, which we assume to be linear. By the Butler-Volmer model, an electrochemical cell can be linearized within a relatively small window of overpotential, and the nonlinear Tafel behavior outside this window is associated with high rate of faradaic reactions [28], which usually should be avoided in neural stimulation. Furthermore, electrochemical impedance spectroscopy (EIS) is only meaningful for electrode-electrolyte interfaces with linear circuit behavior, and as a result, most discrete-element circuits used to model neural electrodes assume linearity. The circuit driver, however, may have nonlinear current-voltage (*i*-*v*) relationship, especially when neither the current nor the voltage output of the circuit driver is directly controlled. For example, for photovoltaic retinal prosthesis, only the light intensity projected onto the photodiodes is directly controlled, and the *i*-*v* relationship of the photodiodes is exponential [23].

#### 2.1.1 Directly controlled driver of current or voltage

For a circuit driven by controlled *I*_*D*_(*t*) or *V* (*t*), we can calculate the steady-state response of the load to periodic stimulus directly from the impedance spectrum. Let impedance *Z*(*ω*) : ℝ ↦ ℂ, as a function of frequency *ω*, be the impedance spectrum of the load, and ***ℱ*** be the Fourier transform. *I*_*D*_, equivalently, *I*_*L*_, and *V* are related by

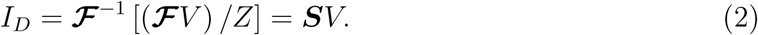

(2) is a linear transform from function *V* to function *I*_*D*_, which we denote with the linear operator ***S***. Since the impedance is typically nonzero and finite, ***S*** is invertible. Therefore, given a stimulus waveform *I*_*D*_(*t*) or *V* (*t*), we can calculate the corresponding response of *V* (*t*) or *I*_*L*_(*t*). [27] proposes a similar approach to study the responses of electrochemical cells in frequency domain, but this method still involves equivalent circuits to calculate the Laplace transform for transient behavior. Our approach avoids the complication of finding an equivalent circuit, and can also be applied to describe the transient behaviour in the non-periodic stimulus, as shown in Section 2.1.3. Note that the DC impedance *Z*(0) is not measured in the standard EIS, and can be determined separately via chronoamperometry or methods alike. However, in neural stimulation, charge should be balanced, and therefore the DC current should be zero.

#### 2.1.2 Nonlinear circuit driver

When neither *I*_*D*_(*t*) nor *V* (*t*) is directly controlled, we can rely on the *i*-*v* relationship of the driver at each time step:

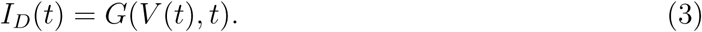

The *i*-*v* relationship can change over time, such as the *i*-*v* curve of an illuminated photodiode, and hence *G* is a function of both *V* and *t*, which may be nonlinear. Knowing *G*, we can study the steady-state response following the principles of HB analysis - comparing the responses of the linear and the nonlinear parts of the circuit, and using optimization tools to minimize the difference [29, 30].

For any given *V* (*t*), the nominal *Ĩ*_*L*_(*t*) is given by (2):

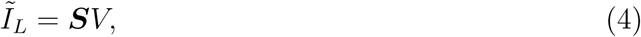

and the nominal *Ĩ*_*D*_(*t*) is given by (3). Note that *Ĩ*_*L*_(*t*) and *Ĩ*_*D*_(*t*) have no physical meanings, because generally *Ĩ*_*L*_(*t*) ≠ *Ĩ*_*D*_(*t*), which violates (1). We can define the difference between *Ĩ*_*L*_ and *Ĩ*_*D*_ as an operator acting on *V* :

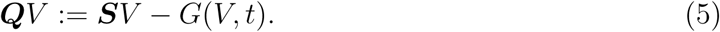

The true voltage response at the port, *V*\*, is obtained if and only if ‖***Q****V* *‖^2^ = 0. We perform optimization on ‖***Q****V*‖^2^ to find *V*\*:

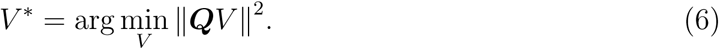

We prove in the Appendix that min_*V*_ ‖***Q****V*‖^2^ = 0 and the solution *V*\*(*t*) is unique, if *G*(*V*) is monotonically non-increasing. The monotonicity is a realistic premise, because higher voltage at the port counteracts the current output.

#### 2.1.3 Transient behavior

HB analysis only provides the steady-state solution of circuits under periodic stimuli [29, 30]. However, the several initial pulses may show transient behavior uncaptured in the steady-state solution, especially when they are monophasic capacitor-coupled. For example, in [23], the initial pulses are not charge-balanced, and the response to them is quite different from the steady state.

The general approach to study the transient behavior of an electrode-electrolyte interface is using the Laplace transform without assuming periodicity, which involves finding its equivalent circuit [27]. Here, we convert the transient behavior into a periodic one by padding the waveform with an additional discharging phase and a checking phase. To calculate the initial pulses between *t* = 0 and *t* = *T*_0_, we insert a discharging phase of length *T*_1_ − *T*_0_ after *T*_0_ and a checking phase of length *T*_2_ − *T*_1_ after *T*_1_, as illustrated in Figure 2.

**Figure 2:**
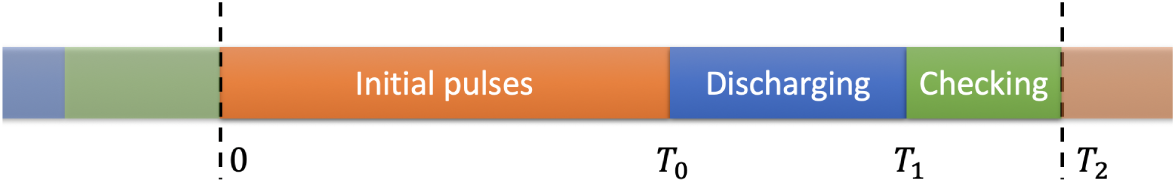
Schematic diagram of the augmented period consisting of the initial pulses, the discharging phase and the checking phase.

In the discharging phase, we keep *V* (*t*) constant at the resting voltage *V*_0_ of the load, and allow any amount of current to flow through the driver. In the checking phase, the circuit driver has the resting *i*-*v* relationship *G*_0_ to verify the load has sufficiently discharged. For the circuit driver, we have

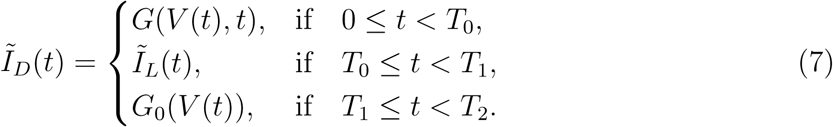

For this scheme to work properly, *T*_1_ − *T*_0_ should be at least several times longer than the time constants of the electrode relaxation and charge redistribution calculated in [31]. To verify the efficacy of the discharging phase, the deviation of *V* (*t*) from *V*_0_ in the checking phase should be negligible compared to its average magnitude between 0 and *T*_0_:

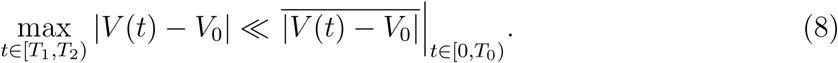

### 2.2 Experimental validation

The circuit of a photovoltaic retinal prosthesis described in [10] can be driven by shining light on photodiodes. However, quantitative analysis of such circuit requires precise measurement of the irradiant power on each photodiode and their light-to-current conversion. To simplify such experiments, we use a current source in parallel with the same photodiode in the dark, which exhibits an identical i-v curve, and therefore allows driving the circuit in an equivalent way as using light.

By Equation (1) of [23], the output current of the photodiode equals the short-circuit photocurrent minus the forward current of the diode under the output voltage. Therefore, the current source and the shaded photodiode *D*_*p*_ combined in parallel are equivalent to the photodiode in light, as shown in the red dashed box in Figure 3. The ON and OFF status of the current source correspond to turning the light on and off, respectively. Because of the limited precision of the current source, we cannot set the OFF current to be strictly zero between pulses. To prevent random drift of *V* (*t*) due to charge accumulation, a Schottky diode *D*_*s*_ is connected in parallel with *D*_*p*_ in opposite polarity, and a negative current that is much smaller in amplitude than the stimulating current is applied between pulses. In the resting state, *V* ≈ −0.1*V*, which is the turn-on voltage of *D*_*s*_. The circuit is designed to function when *V* ≥ 0, and *D*_*s*_ has no effect on the circuit in this range.

**Figure 3:**
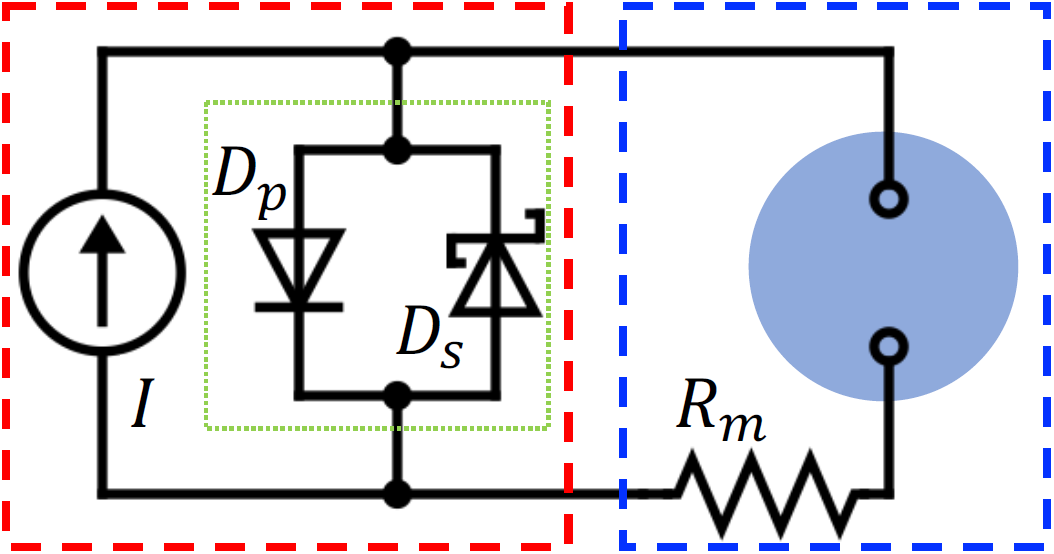
The circuit diagram of the experiment, consisting of the current source, the photodiode *D*_*p*_, the Schottky diode *D*_*s*_, the electrochemical cell (blue circle) and the monitor resistor *R*_*m*_. The circuit driver and the load are noted by the red and the blue dashed boxes, respectively.

We use an electrochemical cell of the 2-electrode setup. The active and the return electrodes are a pair of 80*µm*-diameter platinum disks coated with 400*nm* of SIROF, spaced 250*µm* apart center-to-center on an insulating planar substrate. The electrolyte is a cell-culture medium made of 89% of DMEM/F12 base solution, 10% of fetal bovine serum and 1% of Penicillin(10, 000*U/mL*)-Streptomycin(10*mg/mL*) in volume. The experiment is conducted at 37°*C*. A resistor *Rm* = 5.10*k*Ω is connected in series with the electrochemical cell to monitor the current. The complete circuit is illustrated in Figure 3.

To check linearity of the electrochemical cell, we first measured the impedance spectra at different voltage biases and with different perturbation amplitudes. We then applied Stimulus 1 and Stimulus 2, whose features are summarized in Figure 4 and Table 1, to the current source and recorded the current (*I*_*L*_) and the voltage (*V*) of the load. Next, we used HB analysis to calculate *I*_*L*_ and *V* directly from the impedance spectrum, and compare with the experimental measurements. For comparison, we also fit the discrete-element circuit models to the impedance spectrum and solve the corresponding differential equations for *I*_*L*_ and *V* in time domain.

**Table 1:** Parameters of the periodic waveform output by the current source. *f, τ, i*^+^ and *i*^−^ are defined in Figure 4.

| | $f$ (Hz) | $\tau$ (ms) | $i^+$ ($\mu A$) | $i^-$ (nA) |
| --- | --- | --- | --- | --- |
| **Stimulus 1** | 2 | 50 | 2.679 | -51.2 |
| **Stimulus 2** | 30 | 10 | 9.72 | -255.7 |

**Figure 4:**
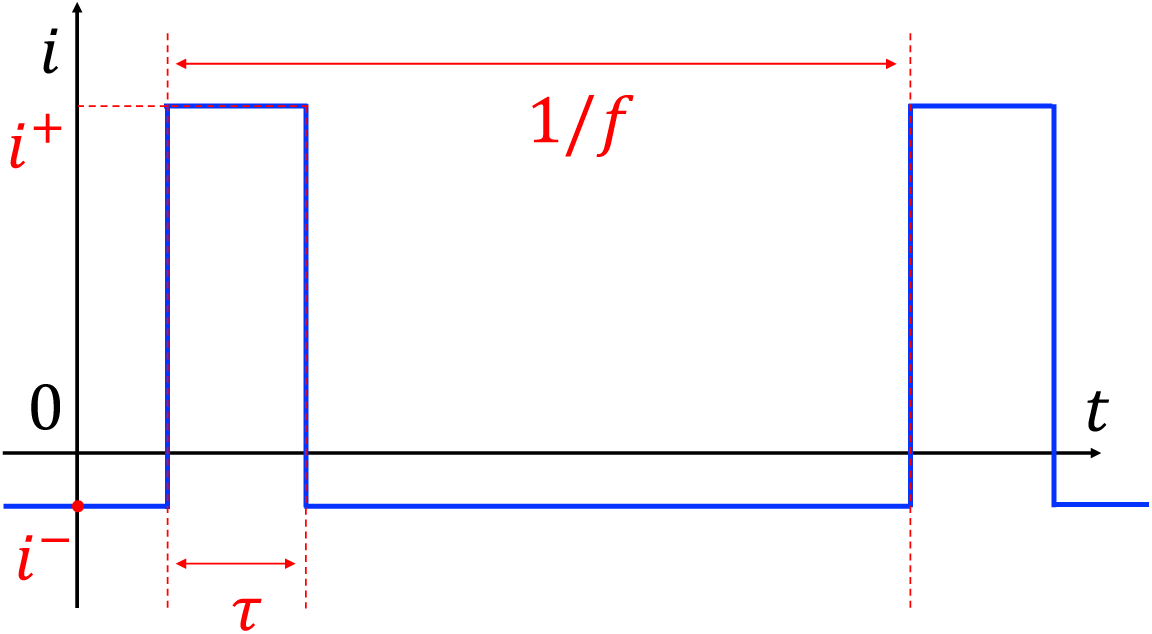
Current output of the current source in Figure 3. Values of the repetition rate *f*, the pulse duration *τ*, the ON current *i*^+^ and the OFF current *i*^−^ are given in Table 1. Plotting is not to scale.

## 3 Results

### 3.1 Validation of linearity by EIS

To validate that the electrochemical cell can be treated as a linear system, we performed EIS at 10 different voltage biases, evenly spaced by 0.1*V* from −0.2*V* to 0.7*V*, with perturbation of 10*mV* root-mean-square (RMS). We also measured the impedance spectrum at 0.3*V* offset with 100*mV* -RMS perturbation. All the spectra, plotted in Figure 5a, match each other very well, confirming linearity of the electrochemical cell. The DC impedance *Z*(0) is determined to be 962*M*Ω by chronoamperometry.

**Figure 5:**
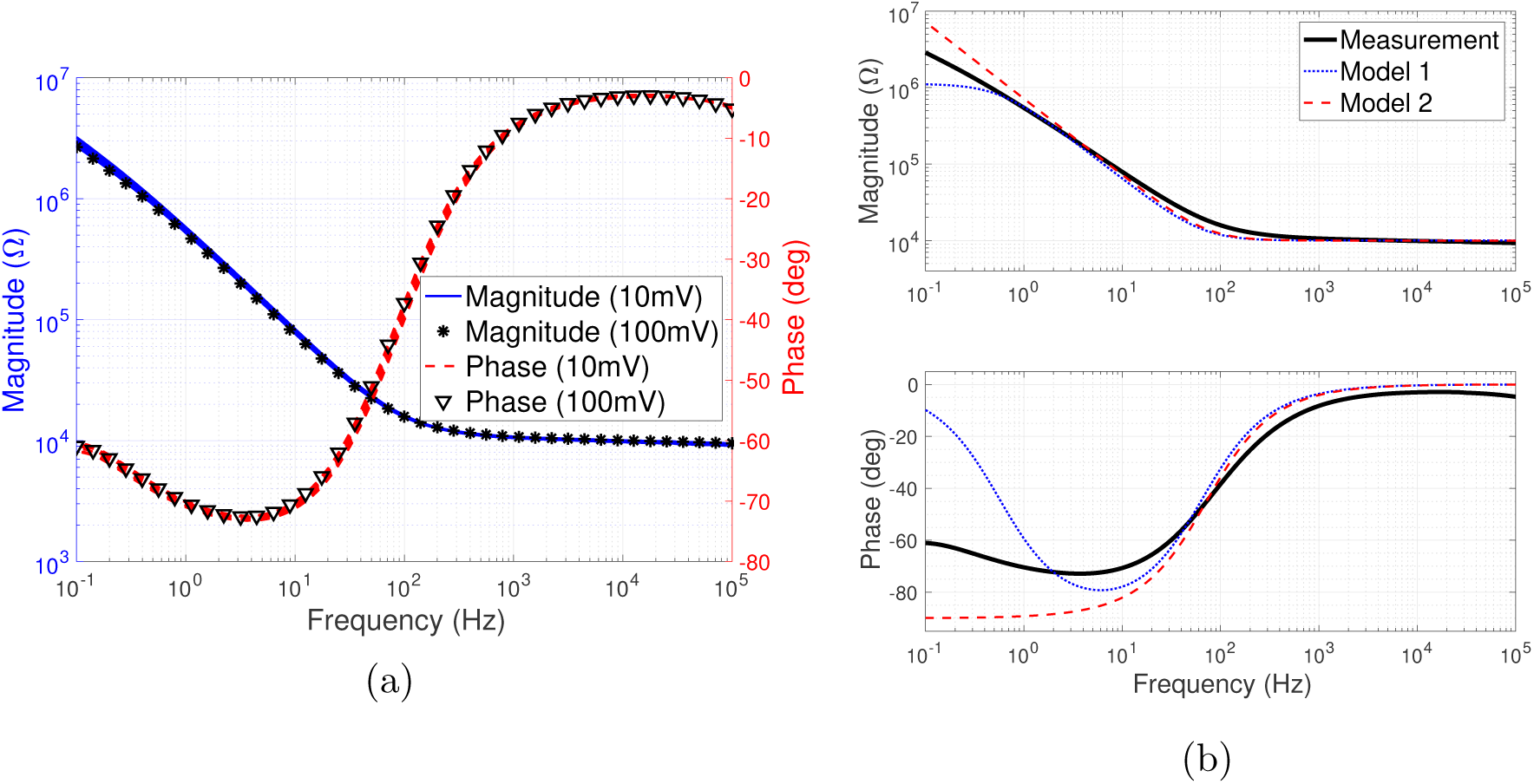
(a) Impedance spectrum of the electrochemical cell, measured at 10 voltage biases evenly spaced between −0.2*V* and 0.7*V*, with 10*mV* -RMS perturbation. An additional measurement is performed at 0.3*V* offset with 100*mV* -RMS perturbation. Each of the colored bundles (solid blue and dashed red) comprises 10 curves, representing the magnitude and the phase, respectively. (b) The impedance spectrum of the electrochemical cell and its two discrete-element approximations shown in Figure 6.

### 3.2 Discrete-element circuit models

A widely used equivalent circuit model of the electrode-electrolyte interface, the Randles circuit, typically includes the constant phase elements (CPEs) [28], the dynamics of which is given in frequency domain. However, since modeling involving a nonlinear circuit driver is more convenient in time domain, a simplification of the Randles circuit, comprising only capacitors and resistors [23, 24, 32] is usually applied. Figure 6a shows one possible simplification consisting of the interface capacitor *C*, the faradaic resistor *R*_*f*_ and the access resistor *R*_*a*_ (Model 1). Sometimes, *R*_*f*_ is neglected and the circuit further simplifies as illustrated in Figure 6b (Model 2) [25]. Generally, we need two sets of lumped circuit to model the working and the counter electrodes, respectively. However, since we use symmetric working and counter electrodes, only one set of the circuit is needed in each model. We use the Levenberg–Marquardt [33] algorithm [ref] to fit the two circuit models to the impedance spectrum in Figure 5a. The spectra of the fit circuits are plotted in Figure 5b, and the values of the discrete elements are shown in Table 2.

**Table 2:**
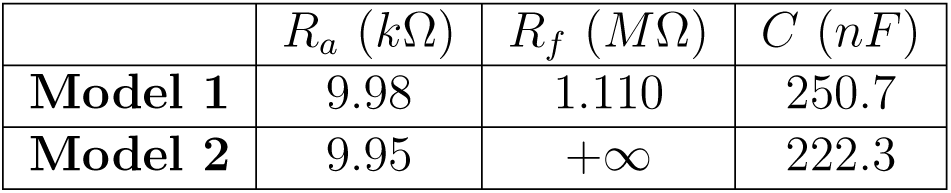
Fit values of the discrete components from the impedance spectrum.

**Figure 6:**
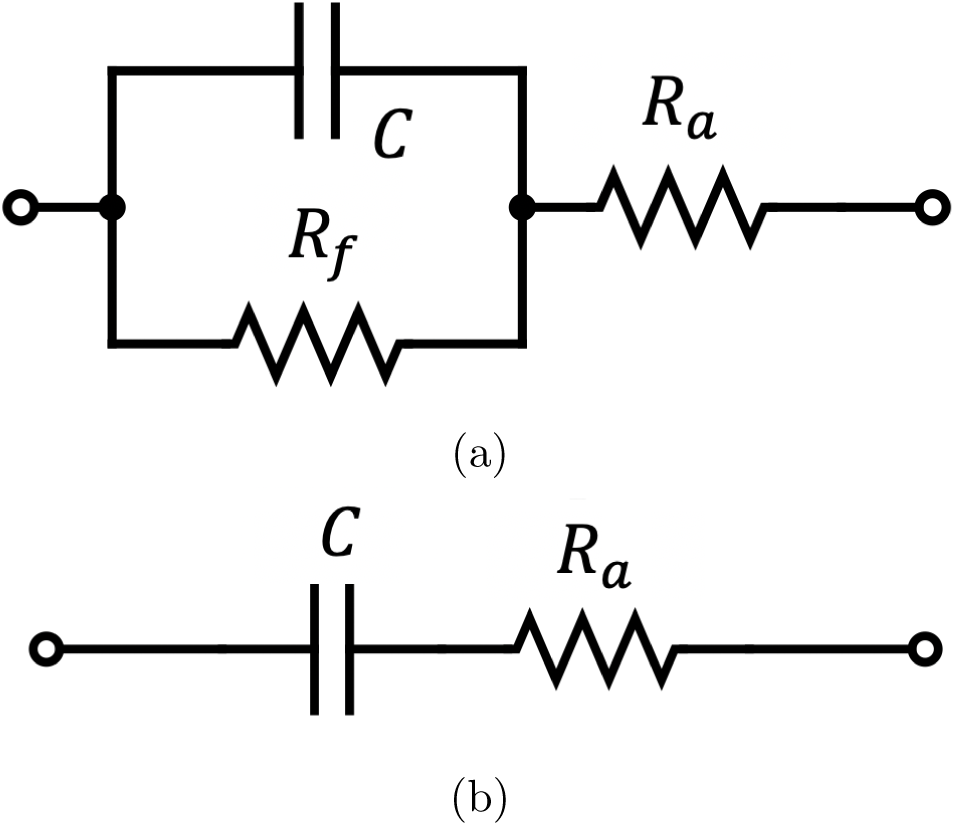
(a) Model 1: simplification of the Randles circuit with the interface capacitance *C*, the faradaic resistor *R*_*f*_ and the access resistance *R*_*a*_. (b) Model 2: further simplification of (a) when *R*_*f*_ is much larger than the total impedance and thereby can be neglected.

The periodic steady-state response to Stimulus 1 with Model 1 and Model 2 can be calculated by solving the differential equations in time domain using Mathematica 11. As shown in Figure 7, the two models produce very different results. Since Model 2 is a special case of Model 1 where *R*_*f*_ is constrained at ∞, Model 1 fits the impedance spectrum better than Model 2. However, the response calculated with Model 2 is more accurate than that with Model 1.

**Figure 7:**
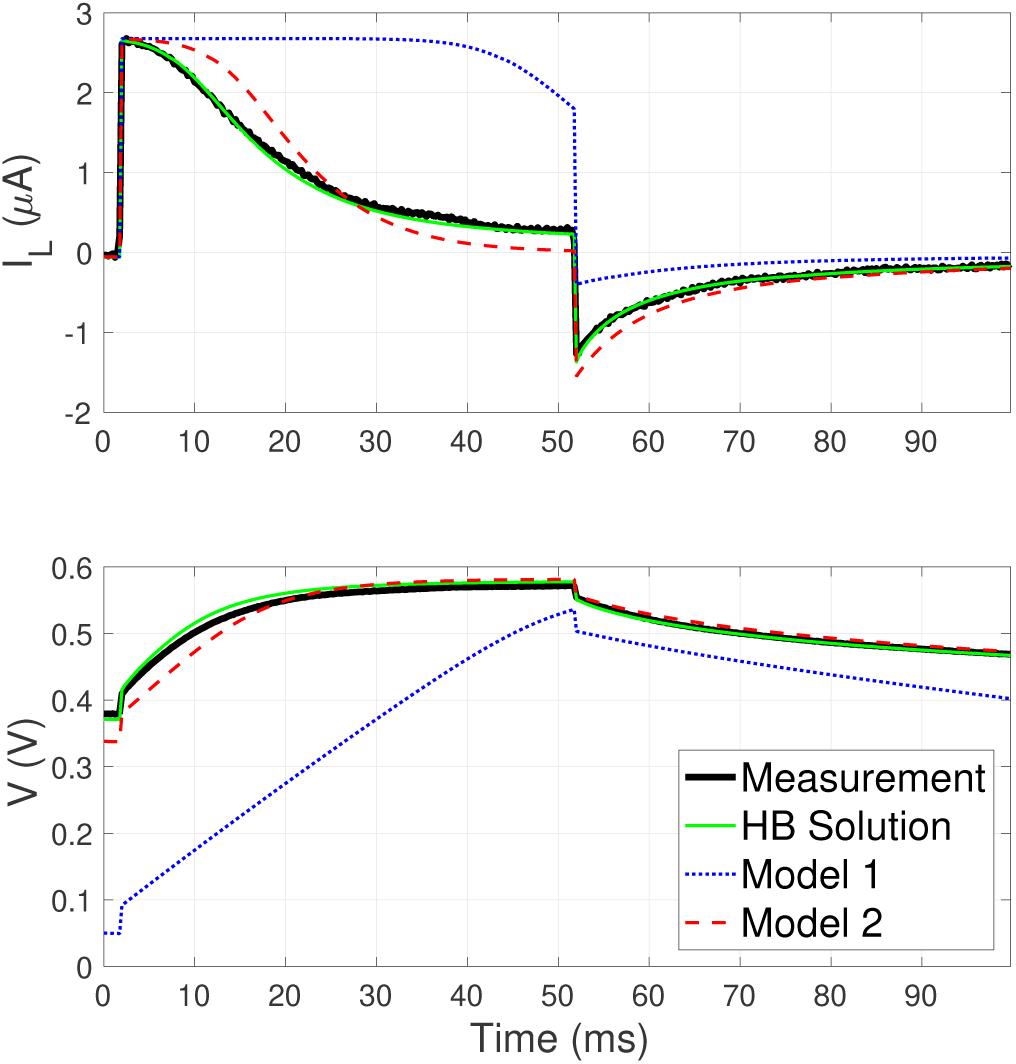
Periodic steady-state response to Stimulus 1. Although Model 1 fits the impedance spectrum better than Model 2, it gives the least accurate prediction of the voltage and current waveforms. The curves are plotted for only 100*ms* to better show the details of the pulse.

### 3.3 HB analysis for steady-state response to periodic stimulus

To find the steady-state circuit response to a periodic stimulus, we implement the optimization in (6) with the built-in nonlinear least square solver (lsqnonlin) in MATLAB (R2017a). As shown in Figure 7, the steady-state response to Stimulus 1 matches the experimental result better than both models in Section 3.2. We also calculated the periodic steady-state response to Stimulus 2 in 50*µs* steps. This optimization took 8 iterations and a few seconds on a laptop computer (3.1*GHz* dual-core CPU, 8*GB* of memory) to converge. Figure 8 depicts the final result plotted with the starting values of *V* (*t*) and *I*_*L*_(*t*), and the two intermediate iterations of the optimization.

**Figure 8:**
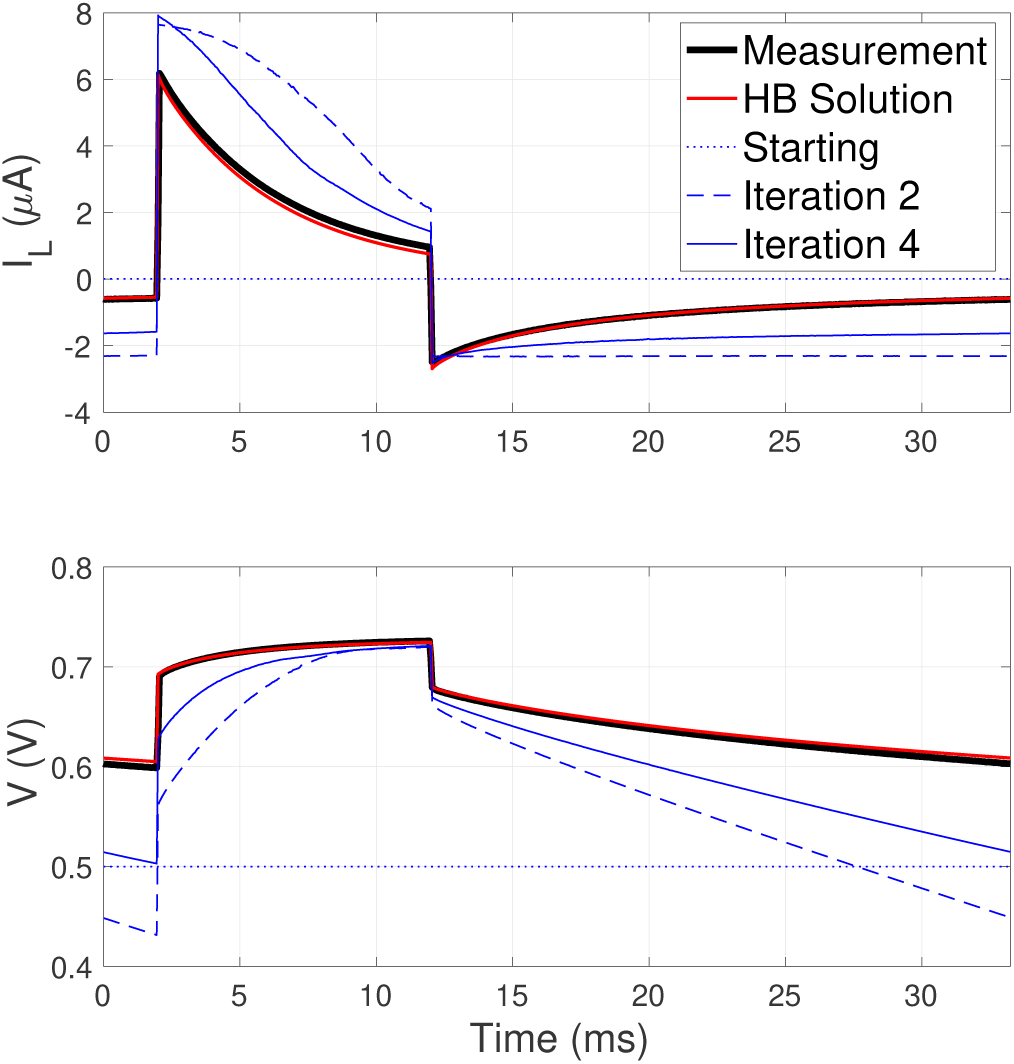
Steady-state response to Stimulus 2. The HB solution converges to the experimental measurement after 8 iterations. The starting value, iteration 2 and iteration 4 are plotted to visualize the optimization process.

### 3.4 HB analysis for transient behavior

Following the scheme outlined in Section 2.1.3, we model the first 115*ms* of the response to Stimulus 2, which includes the four initial pulses. According to Figure 5 in [31], the relaxation time for SIROF electrodes of the same geometry is around 10*ms*. Therefore, we design the discharging and the checking phases to be 60*ms* and 15*ms*, respectively. The transient response based on HB analysis is plotted in Figure 9, and it matches the experimental measurement well. The last pulse in this sequence is already fairly close to the periodic steady-state response to Stimulus 2, shown in Figure 8.

**Figure 9:**
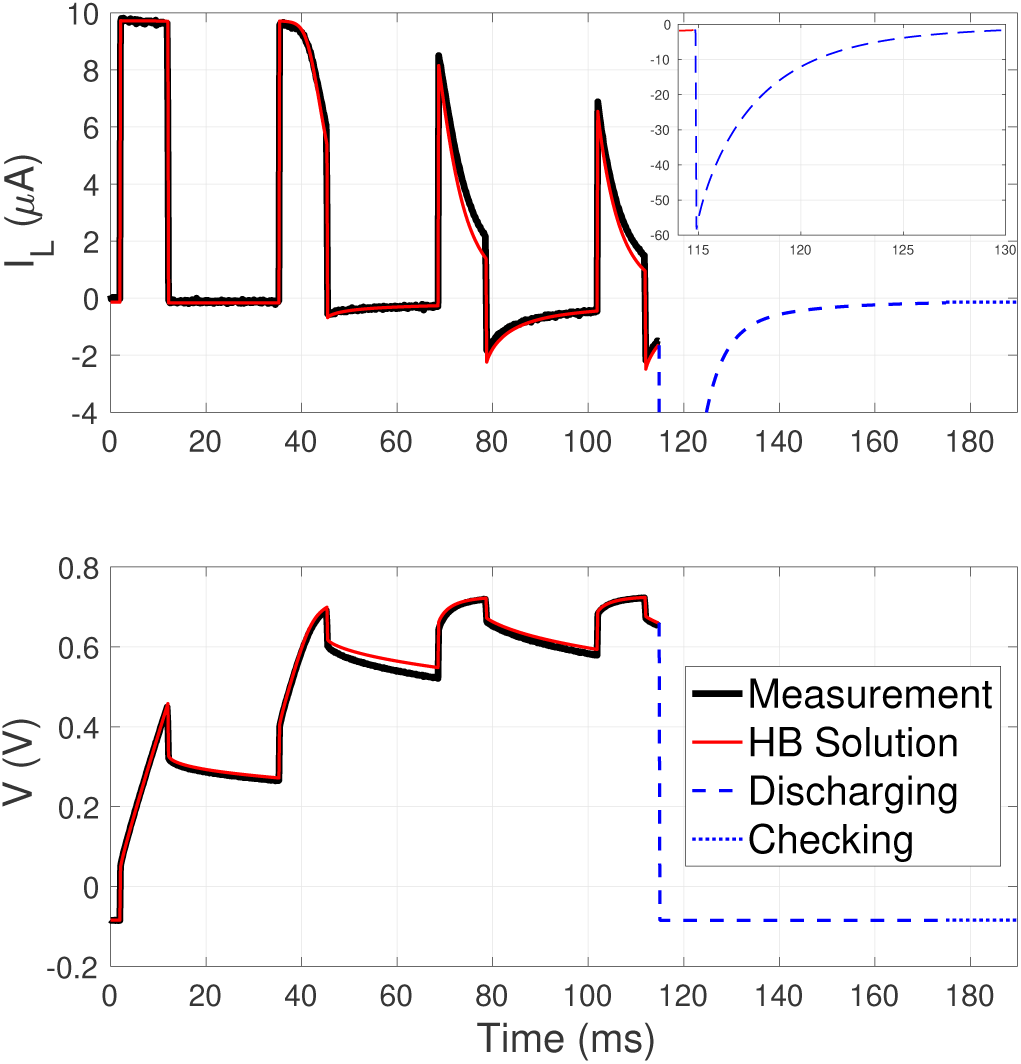
Transient response to Stimulus 2 in the first 115*ms*, featuring the first four pulses. The discharging and the checking phases are 60*ms* and 15*ms*, respectively. The inset of the current (top) panel shows the spike in the discharging phase between 115*ms* and 130*ms*, which is truncated in the main plot.

## 4 Discussion

Successful design of electro-neural interfaces requires comprehensive understanding of the electrode response to given current or voltage stimuli. Only under the assumption that the electrode is ideally polarizable, could the electrode-electrolyte interface be treated as a simple capacitor and CIC has a fixed value. In reality, an electrode material usually exhibits more complex impedance spectrum than what can be represented by a simple *RC* series. One reason for this complexity is that materials capable of storing large amount of charge usually have porous surfaces [19], and some even allow diffusion into the bulk of the electrode [26]. Dynamics of the charge transfer to distributed double layer or redox sites leads to a phase element characteristic of a transmission line in the impedance spectrum [11, 21, 26, 28], and therefore, the CIC becomes waveform-specific. Hence, a computational framework for proper modeling of the electrode-electrolyte interface is nontrivial and important.

One way to model the electrode response is by fitting a discrete-circuit model, comprising capacitors and resistors, to the impedance spectrum, and solving the differential equations in time domain. However, we have demonstrated that the solutions are highly susceptible to small variations in the discrete-element approximations. Figure 5b shows that both discrete-element models approximate the impedance spectrum well in most of the frequency range, except for the low-frequency end. Model 1 includes the faradaic resistor to improve the fitting, but the self-discharge of the electrodes between pulses is overestimated as a result. Calculation with Model 1 shows much higher CIC than in reality. Model 2 does not capture the electrode discharge mechanism, but predicts the response better, even though it approximates the spectrum worse than Model 1. The effect of the faradaic resistor is so prominent because the stimulation period is long compared to the *RC* time constant of the self-discharge loop (*R*_*f*_ and C). The difference between Model 1 and Model 2 should be less significant with stimuli repeated at higher frequencies, illustrating the problem that a match between the discrete-element circuit model and experiment varies with the stimulus shape. A more accurate approximation of the spectrum usually requires CPEs [11, 26–28] and the associated modeling based on the Laplace transform. If the conversion to frequency domain is inevitable, a better approach is to directly apply the impedance spectrum without any approximations. Even though the equivalent circuit models are useful for mechanistic description of various processes in electrochemistry, they may not be the most efficient approach for electro-neural engineering.

Some electro-neural interfaces are driven by nonlinear power sources, such as photodiodes [10] or phototransistors [14]. The proposed HB technique enables computational analysis of circuits with nonlinear elements. By minimizing the difference between the responses of the linear load and the nonlinear circuit driver, the true solution is found if and only if the difference vanishes. HB analysis is adopted because of the convenience to incorporate distributed components described in frequency domain [29]. We show that if the circuit driver has a non-increasing *i*-*v* relationship, the optimization is guaranteed to converge to the unique true solution. Note that our separation of the circuit into a driver and the load is nominal, only for the convenience of discussion. If the load has nonlinear components, it can be lumped with the nonlinear part of the circuit driver to keep the nominal load linear.

The efficient optimization-based HB technique is enabled by advances in optimization research and the growing power of computers. Many optimization algorithms are shown to converge linearly (the distance to the solution decreases exponentially with the number of iterations, i.e., ‖*ϵ*_*n*+1_‖ ≤ *β*‖*ϵ*_*n*+1_‖, where *ϵ*_*n*_ is the distance to the true solution after the *n*^*th*^ iteration and *β* is a constant such that 0 <*β* <1), some even quadratically (‖*ϵ*_*n*+1_‖ ≤ *β*‖*ϵ*_*n*+1_‖^2^) [34]. We used an off-the-shelf optimization tool to demonstrate the baseline capability of HB analysis, whose efficiency can be improved with more sophisticated algorithms. For example, a variety of highly efficient algorithms are developed for convex optimization [35], but our objective functional ‖***Q****V*‖^2^ does not have global convexity - it is convex locally around the solution. One possible approach to utilize the local convexity is to start with fewer sampling points and use non-convex methods to quickly converge to a low-resolution optimum. Then, we can upsample by interpolating the solution, use the interpolated version as the new starting value, and run convex optimization in the neighborhood.

Standard HB analysis can only be applied to periodic waveforms, i.e. it assumes a steady state when repetitive pulsing is applied [29, 30]. We adapted the HB technique to model the transient behavior in the beginning of a long burst of pulses by augmenting the period with the discharging and the checking phases. This adaptation is possible when the electrode-electrolyte interface takes only several cycles before it reaches the steady state, as is the case of typical neural stimulation settings. We demonstrated that the computational workload of the optimization is manageable on a laptop computer, if the number of sampling points in a period is on the order of a few thousands. If the ramp-up would include hundreds or thousands of cycles, as in typical radio-frequency circuits, our adaptation would be computationally inefficient. It also would be inefficient if the pulse duration is very short compared to the period since most sampling points would fall in the inter-pulse interval, the details of which are not of particular interest. Similarly, our adaption is not applicable to the case when the discharge time of the electrode is much longer than the transient behavior.

We developed an experimental technique that enables the testing the characteristics of photovoltaic electro-neural interfaces without using light. The observation that a current source and a shaded photodiode in parallel mimic the *i*-*v* relationship of the illuminated photodiode greatly simplifies the experiments, which would otherwise require optical setup with precise measurements of the irradiant power on the diode and the light-to-current conversion efficiency.

The framework we proposed requires the electrode-electrolyte interface to be linear within the relevant voltage range. Linearity is also the underlying premise of the EIS. We verified the linearity on SIROF electrodes by performing the EIS at different voltage offsets (−0.2*V* to 0.7*V*) and different perturbation amplitudes (10*mV* and 100*mV*). In the future, this method could be extended to cover the case when the impedance spectrum is piece-wise linear, for example, corresponding to different Faradaic reactions within different voltage ranges.

## 5 Conclusions

We established a computational framework to study electro-neural interfaces and the associated electrical circuit dynamics. To model the steady state under periodic stimulus, we directly apply the impedance spectrum to the stimulus in frequency domain. If nonlinear components are involved, we implement HB analysis using a rapidly converging optimization. By adding the discharging and the checking phase in a period, the HB technique is also adapted to study the transient behavior in the beginning of a burst of pulses. We demonstrate that the HB optimization is computationally tractable on a laptop computer, and guaranteed to converge to an unique solution if the circuit driver has a non-increasing *i*-*v* relationship. Comparing the computational results to the experimental measurements, we show superiority of the HB method over the discrete-element circuit approximations.

## Acknowledgements

The authors would like to thank Professor Lenya Ryzhik from Department of Mathematics at Stanford University, Professor Boris Murmann, Dante Muratore and Stephen Weinreich from Department of Electrical Engineering at Stanford University, and Professor Stuart Cogan and Atefeh Ghazavi from Department of Bioengineering at the University of Texas at Dallas for the very helpful discussions. Funding was provided by the National Institutes of Health (Grants No. R01-EY-018608 and No. R01-EY-027786), the Stanford Neurosciences Institute, and Research to Prevent Blindness.

## Appendix

Here we show the existence of a unique solution *V*\* to the optimization (6), such that

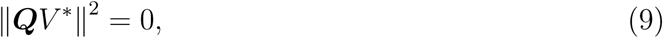

if ‖***Q****V*‖ → +∞ when ‖*V*‖→ +∞, and *G*(*V, t*) is monotonically non-increasing in *V* for all *t.*

*Proof.* First, we show the existence of such *V*\*, by proving

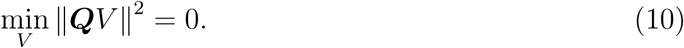

Because ‖***Q****V*‖ → +∞ when ‖*V*‖ → +∞ and ‖***Q****V*‖^2^ ≥ 0, ‖***Q****V*‖^2^ has a minimum. We now show that all extrema of ‖***Q****V*‖^2^ are 0. Suppose ‖***Q****V*‖^2^ is an extremum at *V*_0_. By the Karush–Kuhn–Tucker conditions,

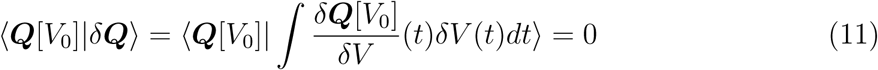

for any variational *δV*. By Equation (5),

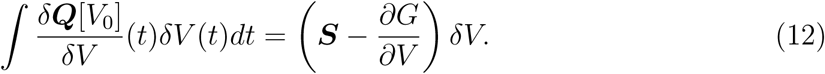

Therefore, we have

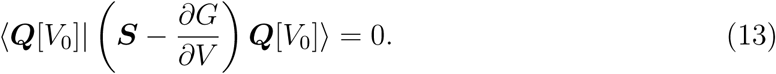

The net power consumption of the load is positive, so by definition, for all *V* ≠ 0, ⟨*V* |***S****V*⟩ > 0. Because 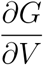 is non-positive, we also have:

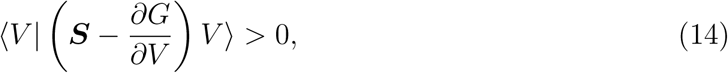

for all *V* ≠ 0, which implies

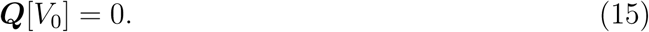

Therefore, ‖***Q****V*‖^2^ has at least one minimum at *V*_0_, and all extrema of ‖***Q****V*‖^2^, including *V*_0_, are 0.

Next, we show the uniqueness of the minimum. We prove, by contradiction, a stronger claim that ***Q****V* is injective, from which the uniqueness directly results. Assume there exist *V*_1_ and *V*_2_, such that ***Q***[*V*_1_] = ***Q***[*V*_2_] but *V*_1_ ≠*V*_2_. We define a function *P* : [0, 1] ↦ ℝ by

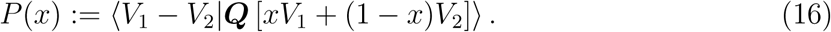

By (14), we have:

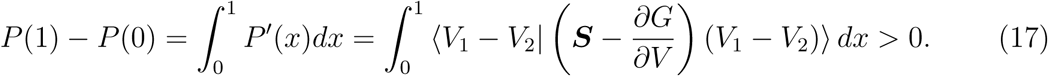

However, we also have

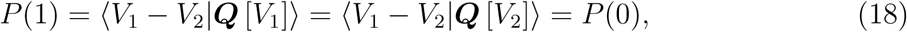

which is a contradiction, and it follows that ***Q****V* is injective.

Therefore, the minimum ‖***Q***[*V*_0_]‖^2^ = 0 is the unique solution *V*\* to the optimization (6).

□

